# Temporal colonization of the gut microbiome in neonatal *Bos taurus* at single nucleotide resolution

**DOI:** 10.1101/2023.03.30.535011

**Authors:** Quanbin Dong, Dongxu Hua, Xiuchao Wang, Yuwen Jiao, Lu Liu, Qiufeng Deng, Tingting Wu, Huayiyang Zou, Luoyang Ding, Shixian Hu, Jing Shi, Yifeng Wang, Haifeng Zhang, Yanhui Sheng, Wei Sun, Yizhao Shen, Liming Tang, Xiangqing Kong, Lianmin Chen

**Affiliations:** Department of Cardiology, The First Affiliated Hospital of Nanjing Medical University, Nanjing Medical University, Nanjing, China; Cardiovascular Research Center, The Affiliated Suzhou Hospital of Nanjing Medical University, Suzhou Municipal Hospital, Gusu School, Nanjing Medical University, Suzhou, China; Changzhou Medical Center, The Affiliated Changzhou No.2 People’s Hospital of Nanjing Medical University, Nanjing Medical University, Changzhou, China; College of Animal Science and Technology, Yangzhou University, Yangzhou, China; Institute of Precision Medicine, The First Affiliated Hospital, Sun Yat-sen University, Guangzhou, Guangdong, China; College of Animal Science and Technology, Hebei Agricultural University, Baoding, China; State Key Laboratory of Reproductive Medicine, Nanjing Medical University, Nanjing, China

**Keywords:** gut microbiome, metagenomics, single nucleotide variations, metabolomics

## Abstract

**Background:** The rumen of neonatal calves is underdeveloped and exhibits limited functionality during early life. Thus, the acquisition and colonization of microbes in the gut are key to establishing a healthy host-microbiome symbiosis for neonatal calves. Microbiome-linked health outcomes appear to be the consequences of individual strains of specific microbes. However, the temporal colonization of pioneering microbial strains and their linkages to the health and growth of neonatal calves are poorly understood.

**Results:** To address this, we longitudinally profiled the gut microbiome of 36 neonatal calves from birth up to 2 months postpartum and carried out microbial transplantation (MT) to reshape their gut microbiome. Genomic reconstruction of deeply sequenced fecal samples resulted in a total of 3,931 metagenomic assembled genomes (MAGs), of which 397 were identified as new species when compared with existing databases of *Bos taurus*. Single nucleotide level metagenomic profiling shows a rapid influx of microbes after birth, followed by strong selection during the first few weeks of life. MT was found to reshape the genetic makeup of 33 MAGs (FDR<0.05), mainly from *Prevotella* and *Bacteroides* species. We further linked over 20 million microbial single nucleotide variations (SNVs) to 736 plasma metabolites, which enabled us to characterize 24 study-wide significant associations (P < 4.4×10^−9^) that identify the potential microbial genetic regulation of host immune and neuro-related metabolites, including glutathione and L-dopa. Our integration analyses further revealed that microbial genetic variations may influence the health status and growth performance of neonatal calves by modulating metabolites via structural regulation of their encoded proteins. For instance, we found that the albumin levels and total antioxidant capacity in neonatal calves were correlated with L-dopa, which was determined by SNVs via structural regulations of metabolic enzymes.

**Conclusions:** The current results indicate that the temporal colonization of microbial strains and MT-induced strain replacement are integral in the development of the gut microbiome of neonatal calves and may help to develop strategies that can improve the health status and growth performance of neonatal calves.

## INTRODUCTION

The colonization of the microbiome in neonatal animals is crucial, as these resident microbes support many functions, including the maturation of the immune system, the utilization and modification of nutrients, and the prevention of pathogen colonization [1, 2]. While dairy cattle harbor a diverse community of ruminal bacteria that provides energy for growth and milk production, the rumen of the neonatal calf is still under development and exhibits limited functionality during early life [3]. Therefore, the acquisition and colonization of the gut microbiome are key to establishing a healthy host-microbiome symbiosis in the neonatal calf. Thus, it’s important to elucidate the temporal dynamics of the gut microbiome during early life to understand the relationships between the gut microbiome and the health status of neonatal calves. Eventually, this will help in designing intervention strategies to achieve better health and growth performance.

Studies in neonatal calves have demonstrated that the microbial taxonomic composition changes dramatically after birth [4–6]. For example, within 24 hours of giving birth, there is a significant increase in the abundances of *Enterobacteriaceae* and *Enterococcus* [4]. After a week of birth, *Lactobacillus* and mucosa-associated *Escherichia* are significantly more abundant [5, 6]. In addition, the population structure of gut microbes in lactating cows and calves differs significantly [7]. Furthermore, the composition and structure of the intestinal microbiome can vary among individuals over time. These observations have laid the foundation for targeted mechanistic investigations into the consequences of host-microbiome crosstalk for the health and growth of newborn calves.

However, investigations into the temporal dynamics of the gut microbiome of newborn calves have mainly focused on the taxonomic and functional composition and have not explored how microbial strains evolve over time. Each microbial species consists of different strains that vary in single nucleotide variation (SNV) and may have different functions [8–10]. Recently, crucial links between microbial genetic composition and longevity [11], as well as multiple disease risk factors [12], have been revealed in humans. Therefore, tracking the temporal dynamics of microbial genetic variations and linking them with the health status and growth of newborn calves may provide us with a new layer of information regarding the role of the gut microbiome in neonatal animals.

In this study, we conducted a longitudinal follow-up of the gut microbiome of 36 neonatal calves from birth up to 2 months postpartum and performed microbial transplantation (MT) to evaluate whether altering microbial strains may influence the health status and growth performance of the calves. We characterized the temporal stability of the gut microbial genomic makeup of the calves, and further investigated the link between temporal variations and the health status and growth of the neonatal animals. To gain additional biological insights, we profiled plasma levels of 736 metabolites at various time points to pinpoint the potential mechanisms behind the microbial genetic impact on the neonatal calves.

## RESULTS

### Genomic assembly resulted in 397 new species-level genomes

The study aimed to investigate the temporal dynamics of gut microbial strains at a single nucleotide resolution and the influence of early microbial intervention on the health and growth performance of neonatal calves. For this, 36 neonatal calves were randomly assigned to three groups: a control group (CON), a rumen microbiota transplantation group (RMT), and an autoclaved rumen fluid transplantation group (RFT). Fecal samples were collected at 15, 35, and 56 days after birth and subjected to metagenomic sequencing using the Illumina NovaSeq-6000 platform. On average, we obtained 36.8 million paired reads for 104 fecal samples collected from newborn calves.

To reconstruct microbial genomes in the gut of neonatal calves, we carried out *de novo* assembly. To maximize the quantity of reconstructed genomes, we followed a single-sample assembly strategy as suggested in a recent study [13]. The detailed workflow is illustrated in Figure 1A. Using this pipeline, we obtained a total of 3,931 metagenomic assembled genomes (MAGs) that met the minimum information about a metagenome-assembled genome (MIMAG) standard [14], with a completeness of ≥50% and <10% contamination. It is noteworthy that the assembled genomes were of high quality, with an average completeness of 84% and a contamination rate of 1.7% (**Table S1**).

**Figure 1.**
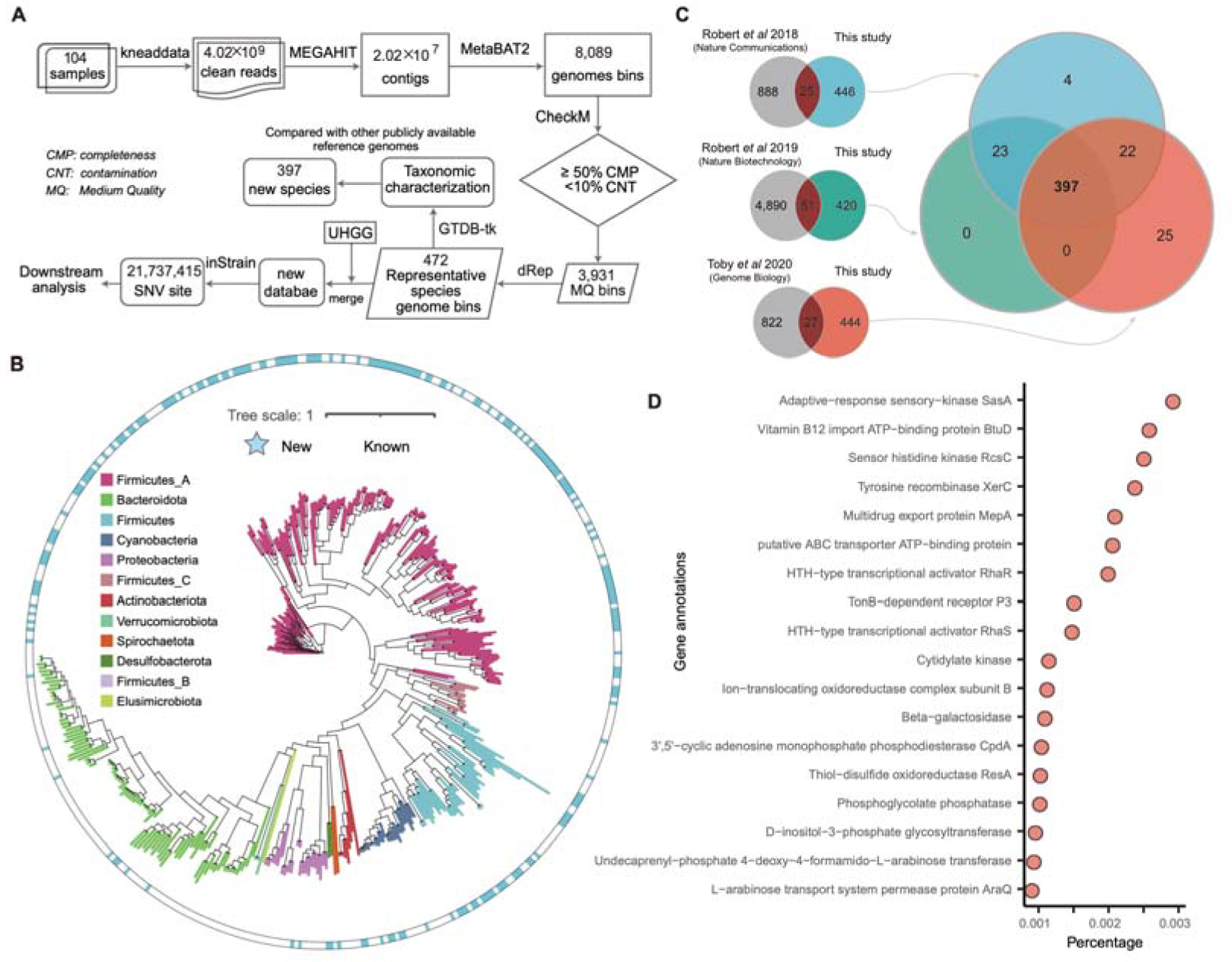
Metagenomic assembled genomes of the present study. **A.** Microbial genomic assembly workflow. **B**. Phylogenetic tree of 472 representative species-level metagenomic assembled genomes (MAGs). The colors on the branches represent the phyla to which the bacteria belong, and the blue color on the outer circle indicates that this is a newly identified species compared with existing genomic references. **C.** Comparison of 472 representative mags with publicly available reference genomes from the three recent studies. **D**. Functional annotation of microbial genes in the 397 newly identified MAGs.

We next checked the taxonomy of those MAGs by clustering all 3,931 genomes with an average nucleotide identity (ANI) threshold of 95%, resulting in 472 representative prokaryotic species (**Table S2**). According to the taxonomic classifications using GTDB-Tk [15], these genomes mainly belong to the phyla *Firmicutes* and *Bacteroidota* (Figure 1B, **Table S2**). We compared these representative genomes with existing microbial genomes obtained from cattle, including RUG [16], RUG2.0 [17] and African MAGs [18]. At 95% ANI, we found that 446, 420, and 444 of them were novel when compared with RUG, RUG2.0, and African MAGs, respectively (Figure 1C). Importantly, 397 out of the 472 representative genomes were not covered in the above-mentioned studies (Figure 1C).

To investigate the functionalities of these novel species, we used Prokka [19] to annotate the functional genes in the newly discovered genomes. Our analysis revealed that these novel species have variable functionalities, ranging from adaptive-response sensory-kinase to glutamine synthetase, while a substantial number of genes were not annotatable, with 47% of the genes being hypothetical (Figure 1D, **Table S3**).

In summary, the newly constructed genomes are a valuable supplement to existing resources for microbiome research in neonatal calves, providing additional insight into the taxonomy and functional capabilities of gut microbes in this population.

### Temporal variability of the gut microbial strains in neonatal calves

Investigating the temporal colonization of gut microbial strains in neonatal calves is crucial as it can provide insights into the mechanisms that drive the development of calves and identify potential targets for interventions to promote host health. Here, by comparing the number of MAGs obtained from each calf at different time points, we found that the number of MAGs obtained in neonatal calves increases over time (Figure S1A). Moreover, this increase was not biased by differences in sequencing depth as significant negative but not positive correlations were observed (Figure S1B), demonstrating a rapid influx of gut microbial strains after birth during the first few weeks of neonatal calf development.

To study the temporal colonization of microbial strains in the gut of neonatal calves, we utilized shotgun metagenomic data to conduct strain-level analysis using single nucleotide variations (SNVs). To achieve this, we combined our MAGs with the UHGG [20] database, which is currently the most comprehensive microbiome sequence resource available. This enabled us to not only investigate study-specific microbial strains but also other existing strains. We aligned sequencing reads to this customized reference database and performed SNV calling using the inStrain [21] algorithms.

In total, 21,737,415 unique SNVs were detected in the 104 samples, derived from 1,209 representative MAGs in our customized reference database, with a range of 440,250 to 1,564,434 SNVs per sample. Similar to the number of assembled MAGs, we observed an increase in the number of SNVs over time across all three groups (Figure 2A), indicating an increase in genomic diversity of the gut microbiome in newborn calves with time. The bacterial species with the highest number of SNVs included *Bacteroides uniformis*, *Parabacteroides faecavium*, and *Phocaeicola vulgatus*. Upon further analysis of within-calf genetic differences in SNVs, we identified MAGs that exhibited significant temporal changes in their SNVs, such as *Alistipes senegalensis*, *Prevotella sp002353825* and *Bacteroides gallinarum*, while *Roseburia inulinivorans*, *Bacteroides intestinalis,* and *Enterenecus faecium* showed relatively low genetic variability (Figure 2B, **Table S4**).

**Figure 2.**
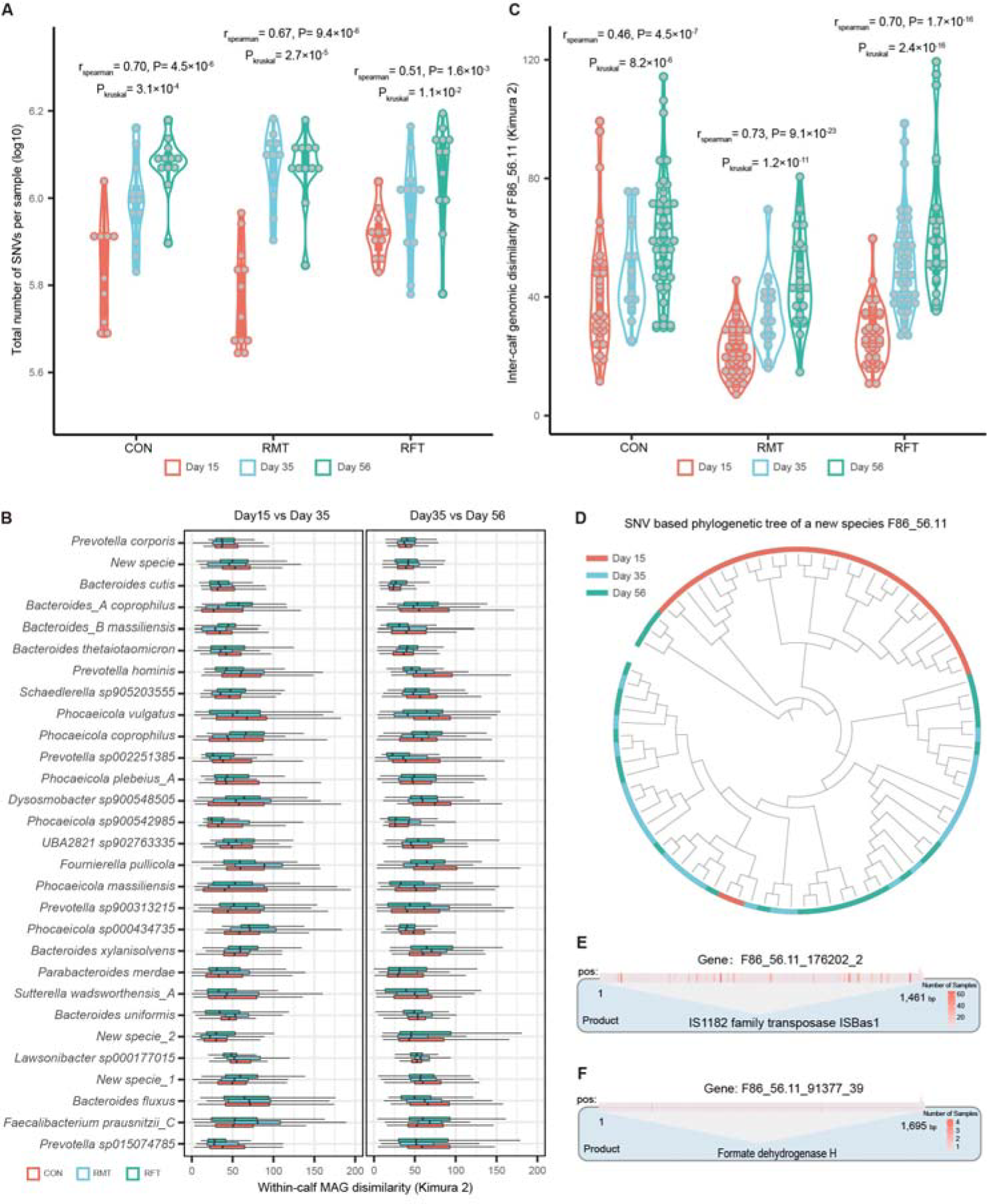
Temporal variability of gut microbial strains in neonatal calves. **A.** The total number of single nucleotide variations (SNVs) detected per sample. Each dote represents one sample. The Spearman and Kruskal tests are used to assess the temporal correlation and within-group differences of SNVs obtained per sample. **B.** The temporal genomic dissimilarity of species-level MAGs is presented. Within-group genomic distance is calculated using the Kimura 2-parameter method. **C.** The inter-calf genomic dissimilarity of a novel species-level MAG *F86_56.11*. The Kruskal test and the Spearman correlation are used to assess temporal differences and associations within the group. **D.** Phylogenetic tree of the novel species *F86_56.11*. Colors represent the time of sampling. **E.** SNVs detected within the genes encoding family transposase are illustrated using a heatmap. Gene length and the number of samples with SNVs are shown. **F.** SNVs detected within the genes encoding formate dehydrogenase H (FDH) are illustrated using a heatmap. Gene length and the number of samples with SNVs are shown.

Interestingly, we found that the genetic variability of a novel species, *F86_56.11*, varied substantially over time. The dissimilarities between calves based on SNVs of this species increased with time and were consistently the lowest on day 15 (Kruskal test, P < 1.0×10^−5^, Figure 2C). Phylogenetic analysis further showed that the species had two distinct strains (Figure 2D), with a significant enrichment of strain 1 observed on day 15 compared to day 35 and day 56 (Fisher exact test, P < 1.07×10^−10^, Figure 2D), indicating the high temporal variability of this bacterium. As the gut environment of neonatal calf changes over time, it places selective pressure on microbes, driving them to evolve and adapt to new conditions, increasing their chances of survival and reproduction.

To investigate the potential mechanisms responsible for the temporal genetic variability observed in *F86_56.11*, we analyzed genes with SNVs that displayed significant temporal differences. Our genomic annotation analysis revealed that several genes that encode family transposase exhibited significant temporal variability in *F86_56.11* (Kruskal test, P < 0.05, Figure 2E, **Table S5**). Transposase is a protein that facilitates the movement of transposons to different locations within the genome. It can bind to transposons and promote their movement within the genome, driving genome mutation [22]. The activity of transposable elements and their associated transposases can significantly impact the evolution and adaptation of bacteria to their environment [22].

In addition, we found that SNVs in a 1,695 bp *fdhF_2* gene that encodes formate dehydrogenase H (FDH) also exhibited temporal differences (Kruskal test, P= 2.5x10^−2^, Figure 2F, **Table S5**). FDH is a crucial enzyme involved in formate metabolism, a key intermediate in the production of short-chain fatty acids (SCFAs) [23]. The temporal changes in FDH could be related to fiber digestion in neonatal calves because acid detergent fiber (ADF) and neutral detergent fiber (NDF) also exhibited significant changes over time (Kruskal test, P < 9.6×10^−4^, Figure S2). These results suggest that the observed temporal genetic variability in microbial strains in neonatal calves could be due to selective pressure on the microbes, which drives them to evolve and adapt to changing conditions in the gut environment. This adaptive process could involve potential mechanisms underlying the temporal changes in transposable elements and the metabolism of SCFAs, such as FDH.

### Microbial transplantation alters the gut microbial strains in neonatal calves

In addition to examining the temporal shifts of gut microbial strains in neonatal calves, we also investigated whether ruminal microbiota transplantation could alter the colonization of specific microbial strains in the gut of these calves. To do this, we compared the genomic dissimilarity (Kimura 2-parameter distance) of 686 MAGs that were present in all three groups, and significant differences were found for 33 MAGs (Kruskal test, FDR < 0.05, **Table S6**). Notably, many of the MAGs that were significantly altered by RMT were from *Prevotella* and *Bacteroides* species, such as *Prevotella mizrahii*, *Prevotella sp900543585*, *Bacteroides caccae*, and *Bacteroides fragilis* (Figure 3A, Figure S3, **Table S6**).

**Figure 3.**
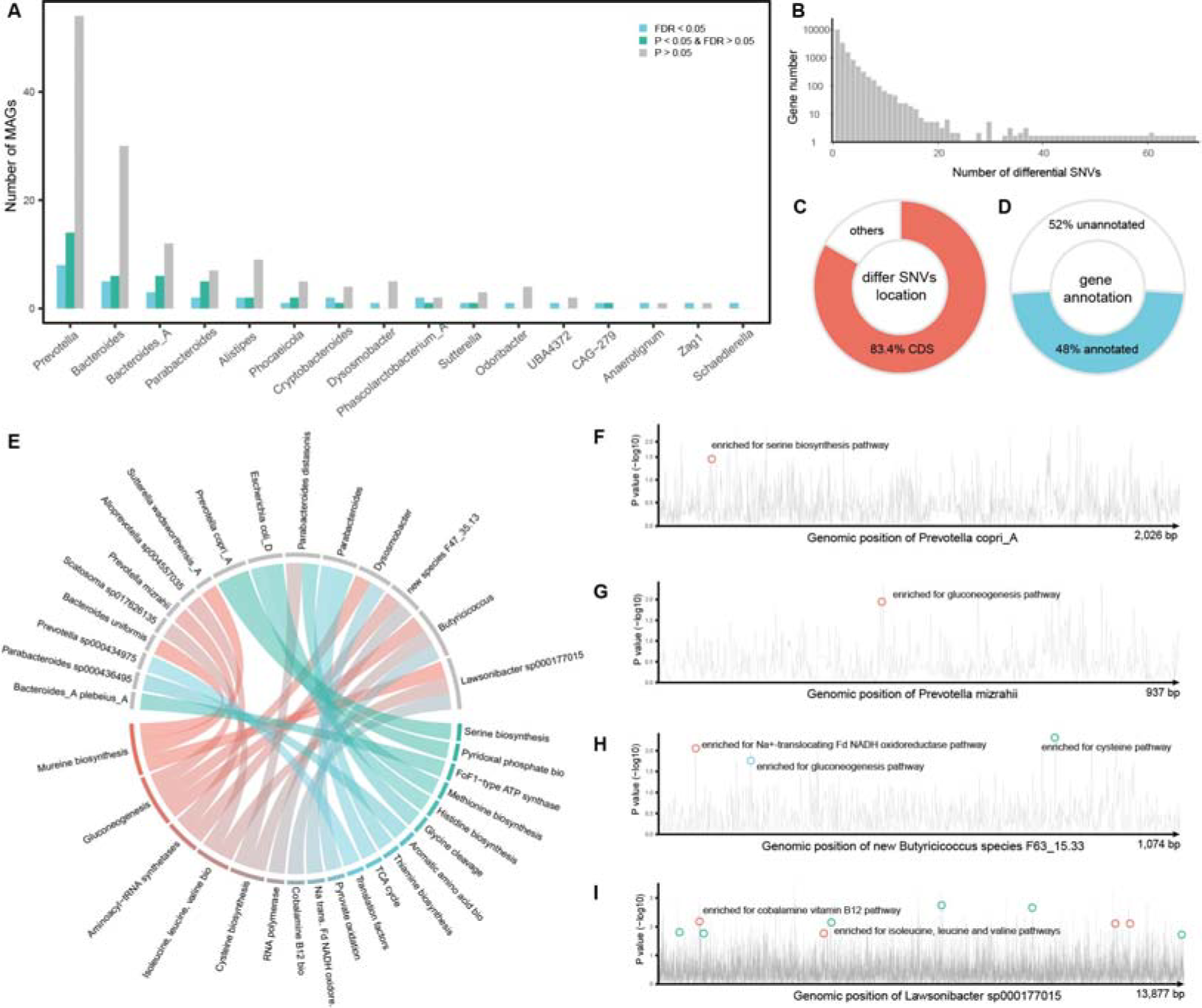
Microbial transplantation alters the gut microbial strains in neonatal calves. **A.** MAGs show differences in their genetic makeup between groups. The X-axis represents different genera, while the Y-axis represents the number of MAGs. The blue bars represent MAGs that showed significant differences at FDR < 0.05. **B.** The distribution of microbial genes with differential SNVs between groups. The X-axis represents the number of differential SNVs within a gene, while the Y-axis represents the number of microbial genes. **C.** The location of differential SNVs within genes. **D.** The percentage of microbial genes that can be functionally annotated. **E**. An overview of 26 significant functional enrichments based on microbial genes with differential SNVs per MAG. **F-I**. Differential genomic loci altered by microbial transplantation and the functional pathway enriched by genes with differential SNVs in *Prevotella copri_A*, *Prevotella mizrahii*, a new *Butyricicoccus species* (*F63_15.33*) and *Lawsonibacter sp000177015*. The X-axis represents the genomic position of SNVs, while the Y-axis displays the P-values (-log10) of differential SNVs between groups.

To further investigate the functional differences underlying the differential MAGs, we analyzed genes with differential SNVs between the groups. In total, we identified 16,497 genes with at least one differential SNV between the groups (**Table S7**), and 3,766 of them (accounting for 22.8% of total genes) had multiple differential SNVs (more than 3, Figure 3B). Notably, 83.4% of differential SNVs were located within the coding sequence (CDS) of genes (Figure 3C, **Table S7**), and 7,914 out of 16,497 genes (48.0%) were functionally annotated (Figure 3D, **Table S7**). These genes encoded a wide range of bioactivities that may have been induced by RMT, including the metabolism of short-chain fatty acids, vitamins, and amino acids (**Table S7**). All of these are important for neonatal calf growth and development, suggesting that early microbial interventions have a significant effect on reshaping the genetic makeup and functionalities of the gut microbiome in neonatal calves.

As we have the COG (clusters of orthologous genes) [24] ids for genes with differential SNVs (**Table S7**), we customized the COG pathway database [24] to calculate pathway enrichment based on genes with differential SNVs per MAG. In total, we identified 26 significant enrichments between 19 microbial pathways and 16 MAGs (Fisher’s exact test, FDR<0.05, Figure 3E, **Table S8**). For example, *Prevotella copri_A* and *Prevotella mizrahii* were enriched for antioxidant-related serine pathways (p=1.4×10^−2^, Figure 3F) and SCFA-related gluconeogenesis pathways (P=1.7×10^−2^, Figure 3G). Moreover, we observed that genes with differential SNVs in a new *Butyricicoccus* species (F63_15.33) were mainly enriched for SCFA-related pathways (Pgluconeogenesis=2.5×10^−2^, PNa+-translocating Fd_NADH oxidoreductase=2.5×10^−2^), as well as antioxidant-related cysteine biosynthesis pathways (p=3.2×10^−2^, Figure 3H). Additionally, *Lawsonibacter sp000177015* was enriched for cobalamin (vitamin B12, P=1.2×10^−5^, Figure 3I) and branched-chain amino acid (BCAA) biosynthesis pathways, including isoleucine, leucine, and valine (P=3.0×10^−3^, Figure 3I). These results further emphasize the importance of microbial interventions during the early days of life in neonatal calves to reshape their gut microbial strains and modulate their metabolism for better health and growth.

### Microbial SNVs associated with plasma metabolites in newborn calves

We characterized significant temporal and RMT-induced differences in the genetic makeup of microbial strains. However, a considerable number of SNVs cannot be functionally annotated (Figure 3D). To gain a deeper understanding of how SNVs might drive host pathophysiology, we hypothesized that metabolites play a crucial role in host-microbe interactions. Therefore, the associations between SNVs and plasma metabolites were assessed. We began by selecting SNVs present in more than 20% of the samples and with a minor allele frequency of 10%. In such a way, 787,964 SNVs were finally associated with 736 untargeted plasma metabolites to identify potential microbial genetic determinants of plasma metabolites in neonatal calves.

A total of 52 significant associations were identified between 50 SNVs from 19 MAGs and 13 metabolites (**Table S9**), with an FDR of less than 0.05 and corresponding P-values less than 4.4×10^−9^. Highly correlated SNVs (r^2^> 0.9, Figure S4) from the same MAGs associated with the same metabolite were filtered, resulting in 24 independent SNV-metabolite associations (Figure 4). The most significant association was found between a SNV from a new species (*F75_56.48*) and the plasma level of phosphatidylcholine (P= 6.63×10^−11^). Among the 24 associations, 8 were observed for SNVs from the genus *Prevotella*, with metabolites mainly including glutathione, D-3-phenyllactic acid, and L-dopa. Glutathione has been reported to affect virulence and bacterial pathogenesis, and the host may use glutathione to modulate its response against bacterial incursions [25]. D-3-phenyllactic acid (PLA) is capable of inhibiting the growth of many microorganisms [26]. Additionally, an SNV in the gene coding agmatine deiminase was associated with dehydroepiandrosterone (DHEA), an important endogenous androgen steroid hormone [27]. Agmatine deiminase is involved in the microbial putrescine biosynthesis pathway [28], which is related to reproductive processes such as spermatogenesis, sperm motility, follicular development, and ovulation [29].

**Figure 4.**
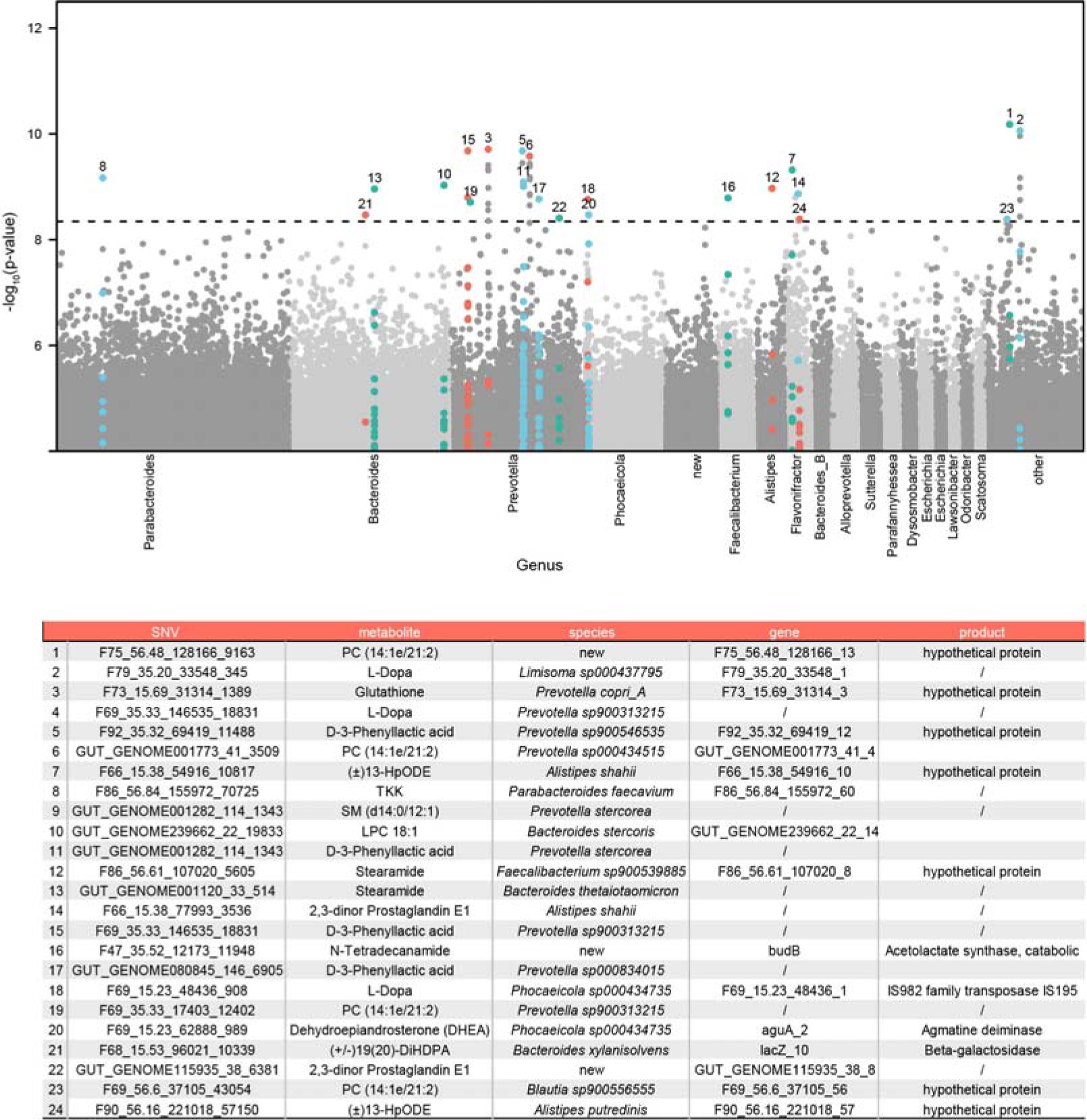
Microbial genetic determinates of plasma metabolites. The Manhattan plot shows the association between microbial single nucleotide variations (SNVs) and 736 untargeted plasma metabolites. Each point in the plot represents a genetic locus, with the X-axis indicating the genomic position of metagenome-assembled genomes (MAGs) in different genera and the Y-axis representing P-values of linear associations. The dashed line represents the study-wise significant line (P < 4.4×10-9), and detailed information about the associations is provided in the table. For SNVs with metabolite association above the study-wise significant line, their associations with other metabolites are also highlighted with the same colors even below the study-wise significant line.

In all, the associations observed between unannotated SNVs and metabolites provide a valuable resource for gaining a deeper understanding of the mechanisms behind host-microbe interactions in neonatal calves. These associations could potentially guide targeted mechanistic investigations to determine the impact of variable microbial strains on the health and growth of neonatal calves. By identifying specific SNVs that are associated with a particular metabolite, we may be able to unravel the complex interplay between the microbial genome and host physiology. Ultimately, this information may lead to the development of new interventions and treatments to improve the health and growth of neonatal calves.

### Microbial SNVs related metabolites associated with phenotypes of neonatal calves

To further understand the relationships between SNV-related metabolites and neonatal calf phenotypes, we performed an association analysis between metabolites and phenotypes. Of the 13 metabolites associated with 50 SNVs (FDR < 0.05, **Table S9**) and the 147 metabolites associated with 14 phenotypes (FDR < 0.05, **Table S10**), we found that 6 metabolites were associated with both (Figure 5A, **Table S11**). The phenotypes involved in these associations included plasma total cholesterol, albumin, malondialdehyde, total antioxidant capacity, as well as the digestibility of ADF (Figure 5A). Notably, most SNV-related metabolites were associated with the total antioxidant capacity of neonatal calves, including L-dopa, D-3-phenyllactic acid, stearamide, glutathione and dehydroepiandrosterone (DHEA). The strongest associations were observed between glutathione and SNV sites at the gene *F73_15.69_31314_3*. For instance, calves with a C or T base at loci F73_15.69_31314_1086 had different levels of plasma glutathione (P= 2.8D×D10^−9^, Figure 5B). Glutathione is an essential molecule for cellular homeostasis and defense against oxidative damage in various diseases [30], and robust correlation between plasma glutathione levels and the antioxidant capacity of neonatal calves was observed (r_Spearman_D=D0.65, PD=D1.2D×10^−6^, Figure 5C). However, the function of this gene based on Prokka was uncharacterized, we then applied AlphaFold2 [31] to predict the structure of the encoded protein (Figure 5B), and the function was further annotated as ATP binding according to the protein structure estimated by DeepFRI [32]. Glutathione is formed by the sequential reaction of L-glutamic acid, L-cysteine, and glycine catalyzed by GSH I and GSH II in the presence of ATP [33]. It was suggested that this protein may bind ATP to influence the synthesis and transport of glutathione [34]. Thus, mutations in this gene may play an important role in the biosynthesis of glutathione via structural regulations (Figure 5D).

**Figure 5.**
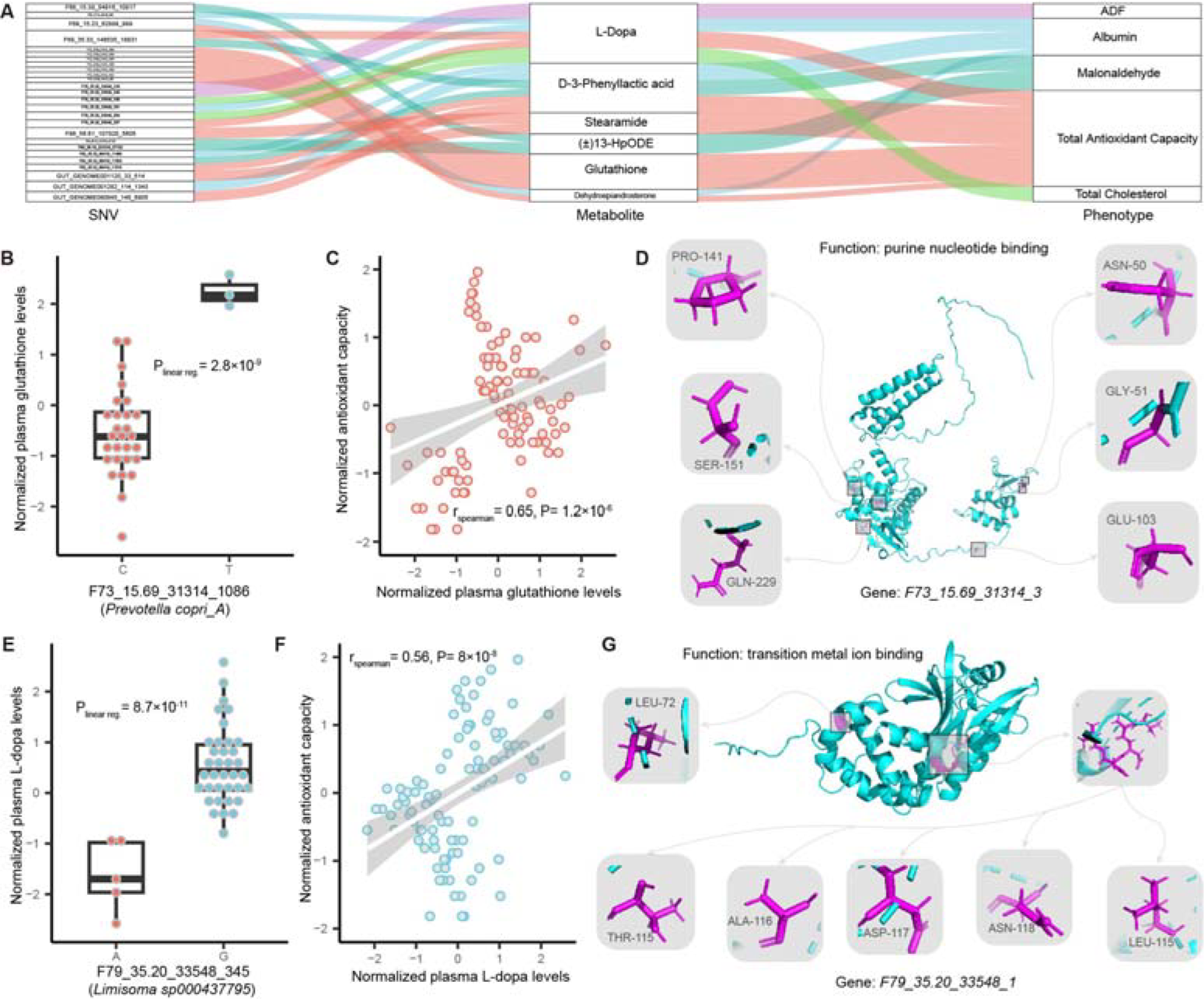
Microbial SNVs related metabolites associated with phenotypes of neonatal calves. **A.** The Sankey diagram shows the association between SNVs, metabolites, and phenotypes of newborn calves. **B.** The association between locus F73_15.69_31314_1086 of *Prevotella copri_A* and the levels of plasma glutathione. **C.** The correlation between plasma glutathione levels and the antioxidant capacity of neonatal calves. The X-axis represents the normalized plasma glutathione levels, while the Y-axis represents the normalized antioxidant capacity of neonatal calves. The fitted linear regression line is shown with a 95% confidence interval. The Spearman correlation coefficient and P-value are shown. **D.** The putative protein structure of gene F73_15.69_31314_3 and the mutation sites on this gene. **E.** The association between different base types at locus F79_35.20_33548_357 of *Limisoma sp000437795* and the levels of plasma L-Dopa. **F.** The correlation between plasma L-Dopa levels and the antioxidant capacity of neonatal calves. The X-axis represents the normalized plasma L-Dopa levels, while the Y-axis represents the normalized antioxidant capacity of neonatal calves. The fitted linear regression line is shown, with the Spearman correlation coefficient and P-value. **G.** The putative protein structure of gene F79_35.20_33548_1 and the mutation sites on this gene.

We also found that multiple SNVs in *F79_35.20_33548_1* were associated with plasma L-dopa levels. Calves with an A or G base at loci F79_35.20_33548_357 had different levels of L-dopa (P= 8.7×10^−11^, Figure 5E), and we observed a significant correlation between plasma L-dopa levels and antioxidant capacity (r_Spearman_D=D0.56, PD=D8.0×10^−8^, Figure 5F). L-dopa is an important precursor for melanin biosynthesis, which is dependent on tyrosinase containing metal ions and plays a vital role in protecting cells from oxidative stress [35, 36]. The function encoded by *F79_35.20_33548_1* is the binding of metal ions, which can be regulated by SNVs through structural changes (Figure 5G). Thus, mutations in multiple sites of the protein F79_35.20_33548_1 may impact the metabolic process of melanin and ultimately affect the antioxidant capacity. Those results provide putative mechanistic insights by identifying specific microbial genetics and functions and highlight which metabolites may be involved in the impact of the gut microbiome on the health of neonatal calves.

## DISCUSSION

In this study, we present a longitudinal investigation of the temporal dynamics of the gut microbiome at the single nucleotide level, and the impact of early microbial interventions on neonatal calves. We assembled gut microbial sequencing data from 104 samples of 36 neonatal calves, resulting in a total of 3,931 metagenomic assembled genomes (MAGs). Our dataset includes 472 unique species-level MAGs (95% ANI), of which 397 have not been previously reported in cattle, thereby providing an additional resource for microbiome research in *bos taurus*. We characterized the temporal dynamics of the gut microbiome at the SNV level, and observed a rapid influx of microbes after birth, followed by strong selection during the first few weeks of life. Additionally, we found that microbial interventions can reshape the gut microbial strains of neonatal calves. We also assessed the association between millions of microbial SNVs and hundreds of plasma metabolites, revealing the genetic regulation of the gut microbiome on host metabolism. Our results show that microbial genetic regulation on host metabolism can be linked to health status and growth performance of neonatal calves.

It is important to understand how the gut microbiome affects the health of neonatal calves by studying the temporal dynamics of the microbiome in early life. Previous studies using 16S rRNA sequencing have examined changes in the composition of the gastrointestinal microbiota in newborn calves during the first few weeks after birth. Our study, which used metagenomic sequencing, provides a more detailed understanding of the genetic makeup of the microbiome over time, revealing that newborn calves are rapidly colonized by microbial strains after birth and then undergo strong genetic selection in the first few weeks of life. We generated thousands of MAGs and identified 397 novel species-level MAGs from fecal samples of newborn calves, which were not previously represented in existing rumen databases. This may be due to the differences between fecal and rumen samples, as well as regional differences. Our study was conducted in China, while other databases have focused on cattle from Scotland [16, 17] or Africa [18]. These newly constructed genomes provide additional resources for future microbiome research in *Bos taurus*.

We investigated the temporal colonization of microbial strains over time in newborn calves by analyzing SNV profiles. We utilized a combination of our sample-specific genomes and the UHGG sequence resource to accurately analyze SNVs in both study-specific and existing microbial strains. Our results showed that the genetic stability of gut microbes varied substantially across different species. Some species, such as *Roseburia inulinivorans*, *Bacteroides intestinalis* and *Enterococcus faecium*, showed relatively low temporal changes over time. Interestingly, previous studies in humans have also shown that some of these species, such as *Bacteroides* [37], are colonized in early life and exhibit high genetic stability in childhood [38]. Moreover, we used metagenomics sequencing to not only identify the strain SNV profiles but also examine the functional differences that may drive these SNV-based genetic clusters. We found that the temporal changes in genes involved in SCFA metabolism may be related to the changes in fiber digestion observed in neonatal calves over time. Our study also demonstrates that metagenomic SNVs are an extra source of information to understand the role of the gut microbiome in neonatal animals. Based on SNVs profile, we found that the gut microbiome of newborn calves can be altered by early microbial interventions at strain level. For instance, the genomic makeup of *Prevotella* and *Bacteroides* species were significantly altered.

In addition, our metagenome-wide microbial SNV association study on 736 plasma metabolites identified 24 independent SNV-metabolite associations. One third of these associations were observed for SNVs from the genus *Prevotella* and the related metabolites, including glutathione, D-3-phenyllactic acid, and L-dopa. Notably, some of these metabolites were also associated with phenotypes of neonatal calves. By identifying the genetic determinants of plasma metabolites in neonatal calves, our study sheds light on the potential role of SNVs in driving host pathophysiology. Specifically, the significant associations between SNVs from microbial strains and plasma metabolites suggest that microbial genetic variation may play a crucial role in shaping host-microbe interactions and contribute to the regulation of host metabolism. These findings highlight the importance of further investigating the molecular mechanisms underlying the observed associations and their potential implications for understanding the interplay between host and microbial genetic variation in health and disease.

The study presented in this paper provides valuable insights into the temporal dynamics of the gut microbiome in neonatal calves and their potential linkages to their health and growth. However, there are several limitations that need to be acknowledged. Firstly, we have characterized many novel microbial strains and have highlighted their importance in neonatal calves. However, the ability to culture and isolate those strains is currently unknown, and further work is needed to determine their viability and potential for experimental validation. Secondly, it is important to note that associations observed in this study do not necessarily imply causation. Additional research is needed to investigate the mechanisms underlying the observed associations and their potential implications for improving early life gut microbiota in neonatal calves. Despite these limitations, the study provides novel insights into the temporal dynamics of the gut microbiome in neonatal calves and highlights the potential for future research to develop strategies for improving gut microbiota in early life.

---

**Figure S1.**
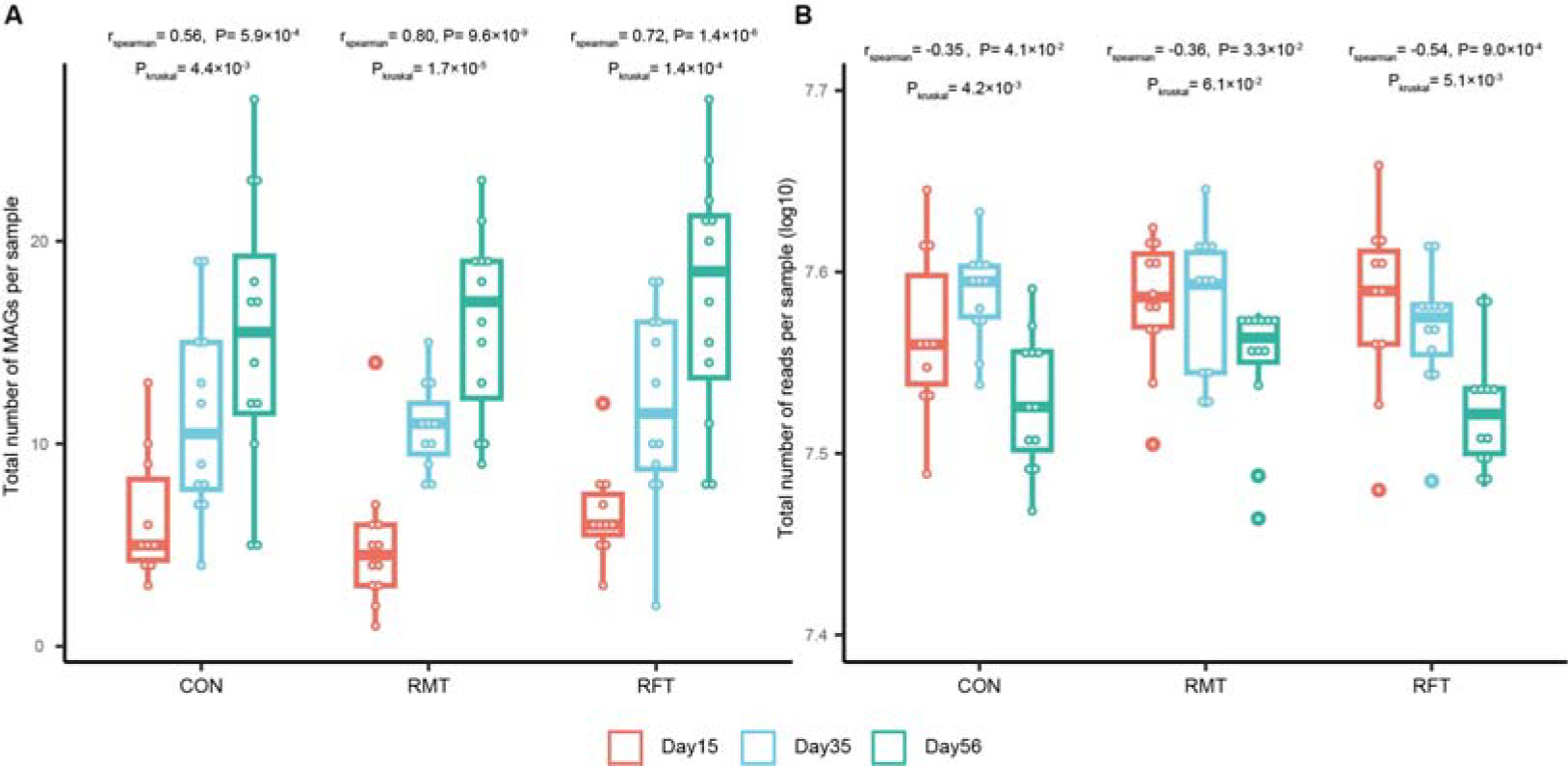
Temporal differences in MAGs and sequencing depth. **A**. Temporal changes of MAGs obtained per sample. Spearman and Kruskal tests are used to assess the temporal correlation and within-group differences of MAGs obtained per sample. **B**. Temporal changes of sequencing depth per sample. Spearman and Kruskal tests are used to assess the temporal correlation and within-group differences of sequencing reads obtained per sample.

**Figure S2.**
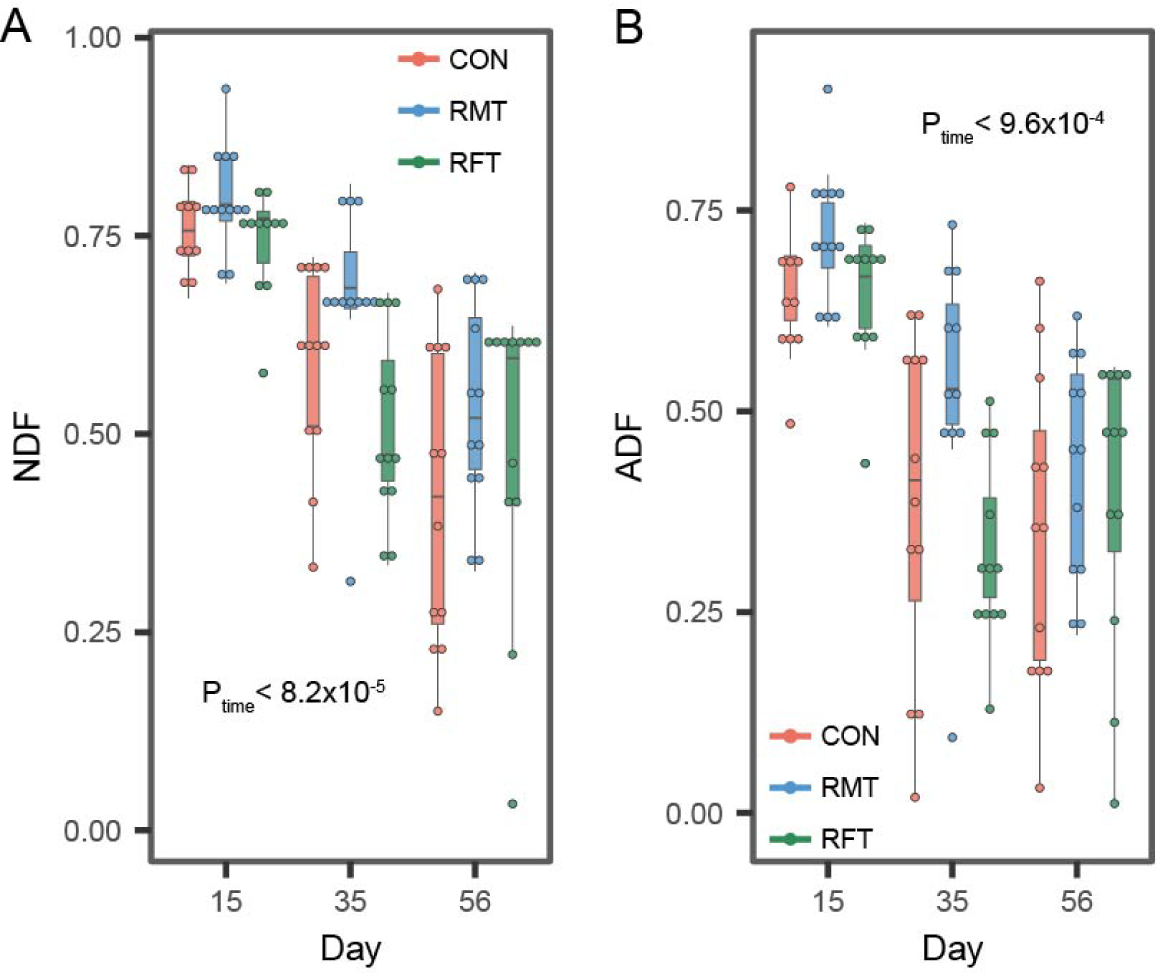
Temporal changes of fiber digestibility in neonatal calves. **A.** Temporal changes of neutral detergent fiber (NDF) digestibility. Each dot represents one sample. **B.** Temporal changes of acid detergent fiber (ADF) digestibility. The P-values from the Kruskal tests are shown. Each dot represents one sample.

**Figure S3.**
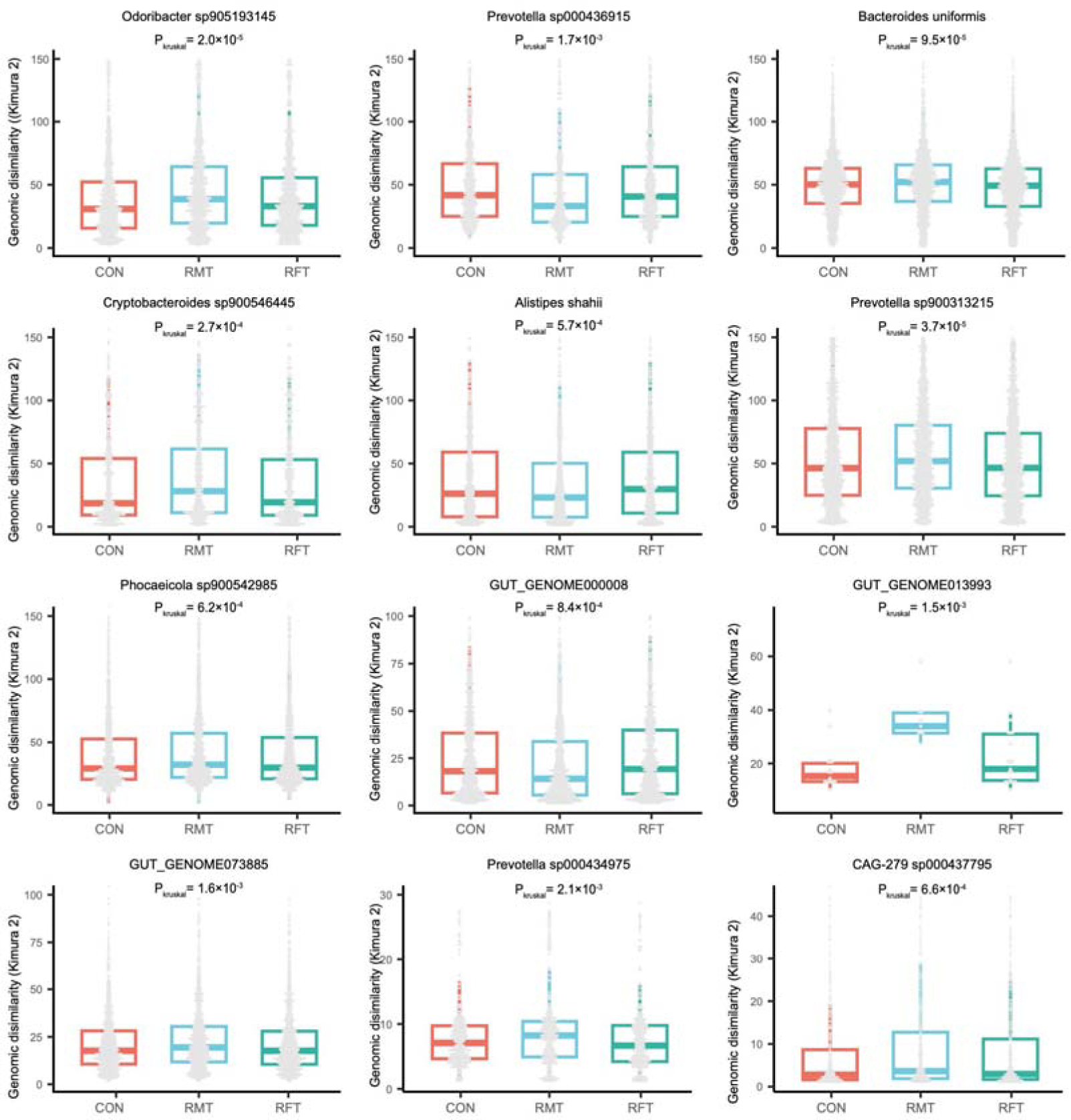
Genomic dissimilarity of MAGs between groups. Inter-calf MAG dissimilarities are calculated using the Kimura 2-parameter method. The Kruskal test is used to assess differences between groups.

**Figure S4.**
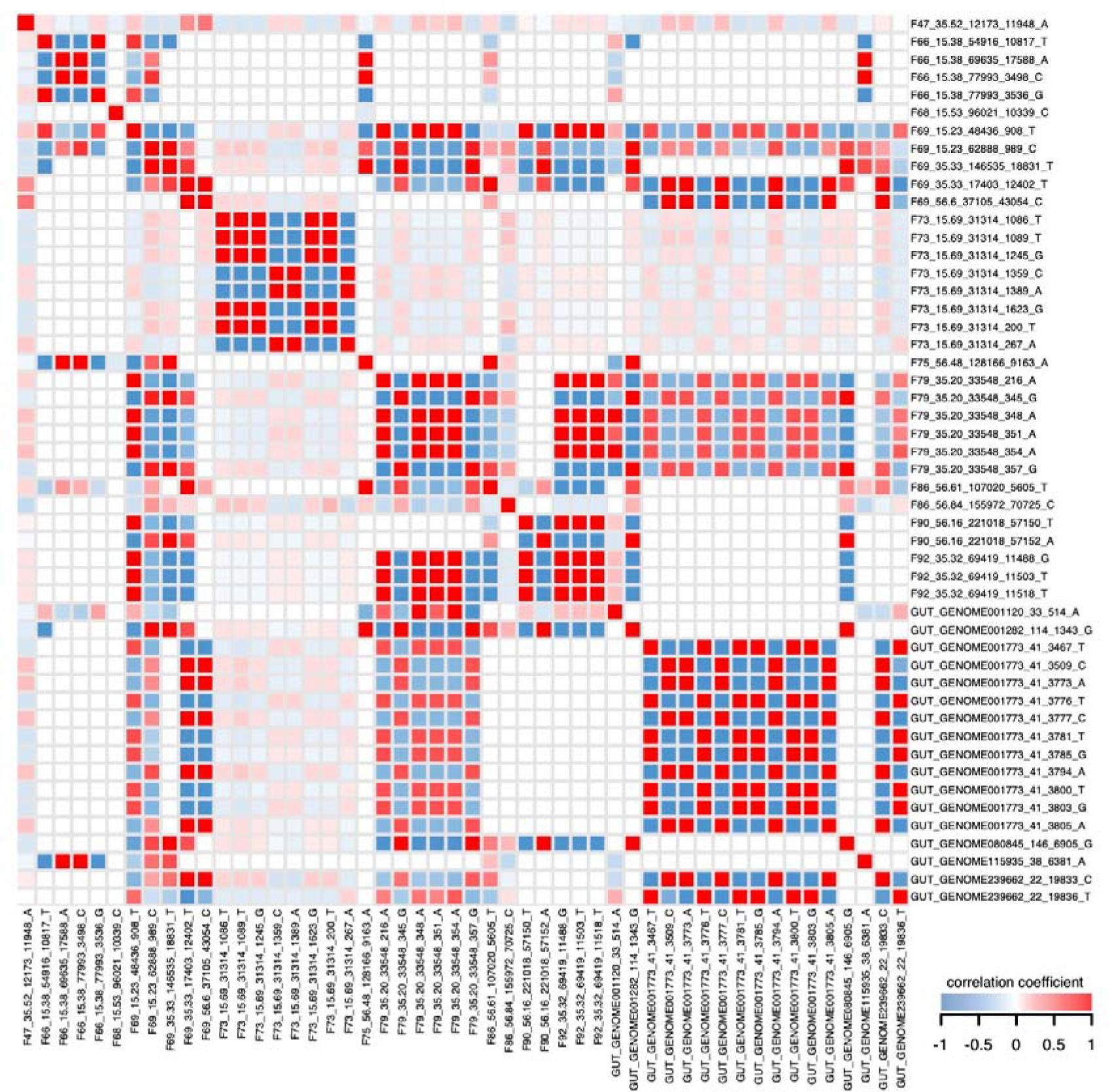
Inter-correlations of microbial SNVs. The heatmap shows Spearman correlations between SNVs with significant associations to plasma metabolites. The darkness of colors represent the correlation strength.

## TABLE LEGENDS

**Table S1.** Quality characteristics of the 3,931 reconstructed microbial genomes

**Table S2.** Taxonomic classification of the 472 representative species-level genomes

**Table S3.** Genomic annotation of the 397 newly reported species-level MAGs

**Table S4.** Within-calf genomic dissimilarity of MAGs at different time intervals

**Table S5.** Genes with SNVs displaying significant temporal differences in *F86_56.11*

**Table S6.** Genomic differences of individual MAGs between groups

**Table S7.** Annotation of the 16,497 genes with differential SNVs between groups across different time points

**Table S8.** Microbial pathway (COG) enrichments based on genes with differential SNVs

**Table S9.** Association between SNVs and plasma metabolites in newborn calves

**Table S10.** Correlation between plasma metabolites and phenotypes of newborn calves

**Table S11.** Metabolites associated with MAGs affected by RMT also associated with phenotype of newborn calves

## METHODS

### Animals

In this study, 36 newborn calves were randomly assigned to three groups and followed for two months after birth. The groups included a control group (CON), a rumen microbiota transplantation group (RMT), and a rumen fluid transplantation group (RFT). The newborn calves were trained to feed milk using a bucket and then transferred to individual calf hutches. Starter was provided *ad libitum* three days after birth and once daily in the morning thereafter. Pasteurized whole milk was fed twice daily at 0800 and 1800 h using a bucket, and the calves were weaned 56 days after birth. RMT and RFT were performed by veterinarians, where the ruminal fluid used in RMT and RFT was collected from a healthy cattle (4-year-old, 600kg, in the dry period,) with a permanent rumen cannula 2 hours after the morning feed. Fresh ruminal fluid was mixed with raw milk and fed to the calves in the RMT group immediately after collection. For the RFT group, the ruminal fluid was autoclaved before feeding. A volume of 50 mL, 80 mL, and 110 mL of ruminal fluid was fed from day 7 to day 11, day 21 to day 25, and day 42 to day 46, respectively. Fecal and blood samples were collected at 15, 35, and 56 days after birth.

### Blood biomarkers

Plasma samples were used to analyze the concentrations of blood urea nitrogen (BUN), glucose, total cholesterol and triglycerides, while serum samples were used to analyze the concentrations of total protein, albumin, alkaline phosphatase, aspartate aminotransferase (AST), alanine aminotransferase (ALT), total antioxidant capacity and malondialdehyde. The concentrations of blood biomarkers were analyzed using commercial kits from Nanjing Jiancheng Bioengineering Institute (Nanjing, China).

For the un-targeted metabolome analysis, plasma samples were resuspended with prechilled 80% methanol. The samples were incubated on ice for 5 minutes and centrifuged at 15,000 g, 4°C for 20 minutes. The resulting supernatant was injected into an LC-MS/MS system consisting of a ThermoFisher Vanquish UHPLC system coupled with an Orbitrap Q ExactiveTMHF mass spectrometer. Raw data files generated by UHPLC-MS/MS were processed using Compound Discoverer 3.1 (CD3.1, ThermoFisher) to perform peak alignment, peak picking, and quantitation for each metabolite. The normalized data was used to predict the molecular formula based on additive ions, molecular ion peaks, and fragment ions. Peaks were then matched with the mzCloud (https://www.mzcloud.org/), mzVault, and MassList databases to obtain accurate qualitative and relative quantitative results. Metabolite annotations were performed using the KEGG database (https://www.genome.jp/kegg/pathway.html), HMDB database (https://hmdb.ca/metabolites), and LIPIDMaps database (http://www.lipidmaps.org/).

### Digestibility

To determine the digestibility of the feed, acid detergent insoluble ash was used as an internal marker [39]. Fecal, starter, and milk samples were collected from each calf and pooled from Day 18 to Day 20, Day 33 to Day 35, and Day 54 to Day 56. The pooled samples were dried at 55 °C for 48 h, ground through a 1-mm screen, and analyzed for dry matter (DM, method 930.15) and crude protein (CP, method 996.11) content according to AOAC International. The contents of neutral detergent fiber (NDF) and acid detergent fiber (ADF) in the starter and feces were measured using heat stable α-amylase and sodium sulfite as described by Van Soest et al. [40]. The apparent total tract digestibility was estimated according to the method of Rice et al. [41].

### Metagenomic data generation and preprocessing

Fecal samples were collected from newborn calves and promptly placed in a freezer (-20°C) within 15 minutes of production. The samples were then transported to the laboratory on dry ice and stored at −80°C until further processing. Fecal DNA was isolated using the QIAamp Fast DNA Stool Mini Kit (Qiagen, cat.51604), and sequencing was performed using the Illumina NovaSeq-6000 platform. The sequencing facility discarded low-quality reads from the raw metagenomic sequencing data, and Bowtie2 (v.2.1.0) was used to remove contamination reads [42, 43]. After filtering, an average of 36.8 million (sd = 3.6 million) paired reads per sample were obtained for subsequent analysis.

### *De novo* assembly and binning

To reconstruct metagenome-assembled genomes (MAGs) from the fecal samples, a bioinformatics pipeline was used as shown in Figure 1A. After removing low-quality and contamination reads, adapter trimming was performed using Trimmomatic (v.0.33) [44], and the resulting clean reads were used as input for MEGAHIT (v.1.2.9) [45]. This resulted in 20,149,780 contigs longer than 200 bp from a total of 2,673,254,051 reads. Reads were then mapped back to the filtered assembly using Bowtie2 (v.2.1.0), and the resulting BAM files were converted to BAM format using SAMtools (v.1.15) [46]. Coverage was calculated using the jgi_summarize_bam_contig_depths script from the MetaBAT2 (v.2.12.1) package [47], and contig binning was performed using MetaBAT2 with contigs shorter than 1.5 kb discarded.

### Genome quality

After binning, we used CheckM (v.1.1.3) [48], which is based on the copy number of lineage-specific single-copy genes, to estimate the quality (completeness and contamination) using the ‘lineage_wf’ workflow. We selected only genomes that passed the following criteria: ≥50% genome completeness and <10% contamination. This also meets the “medium-quality draft” criteria according to recent guidelines [14]. After filtering through this process, a total of 3,931 reconstructed genomes were identified, and the quality statistics, which were measured by analyzing single-copy core genes, are shown in **Table S1**.

### Species-level representative MAGs

To cluster the total set of 3,931 genomes at an estimated species level, we used dRep (v.3.2.2) [49]. We extracted the MAGs displaying the best quality and representing individual metagenomic species. dRep was run with options -pa 0.9 (primary cluster at 90%), -sa 0.95 (secondary cluster at 95%), -cm larger (coverage method: larger), and -con 10 (contamination threshold of 10%). The 95% threshold is commonly used for species-level clustering [50]. Genomes were scored based on their completeness, contamination, genome size, and contig N50, with only the highest-scoring MAG from each secondary cluster being retained as the winning genome in the dereplicated set. Finally, we selected 472 species-level representative MAGs.

### Taxonomic classification and phylogenetic analyses

Taxonomic annotation of each reconstructed genome was performed using GTDB-Tk (v.2.0.0, database release 207) [15, 51] and the ‘classify_wf’ function with default parameters. GTDB-Tk proposes bacterial taxonomy through the concatenation of 120 ubiquitous single-copy proteins, and the producing sequence alignments were used to generate a maximum-likelihood tree. The tree was visualized and annotated using Interactive Tree Of Life (iTOL, v.4.4.2) [52].

### Gene prediction

The coding sequences (CDS) for each of the 3,931 MAGs were predicted and annotated using Prokka (v1.13.3) [19], which utilizes Prodigal (v2.6.3) [53] with the following options: ‘-c’ (predict proteins with closed ends only), ‘-m’ (prevent genes from being built across stretches of sequence marked as Ns), and ‘-p single’ (single mode for genome assemblies containing a single species).

### Single nucleotide variant calling

To improve accuracy and comprehensiveness, we combined our study-specific genomes and a public genome database (UHGG) to build the reference genomes. First, we merged all 472 species-level genomes and the entire UHGG genome collection, which contains all microbial species known to exist in the human gut so far, into a single FASTA file and created a Bowtie2 mapping index from it. The resulting FASTA file contained all the genomes we wanted to profile. Second, we created a scaffold-to-bin file that lists the genome assignment of each scaffold to show which scaffolds came from which genomes using the parse_stb.py script that comes with the program dRep (v.3.2.2). Subsequently, we mapped our metagenomic reads to the well-built reference database using Bowtie2 (v.2.1.0) to create BAM files. Next, we predicted genes for each genome using Prodigal (v.2.6.3) to create a genes file for gene-level profiling. Once all the necessary files were prepared, we called single nucleotide variants (SNVs) by running inStrain profile (v.1.3.2) with default parameters for each sample. We only selected SNVs whose “class” type is “SNV” as defined by inStrain and combined the scaffold, position, and reference base into a single site.

### Genomic dissimilarity

To compare the genomic differences between samples, we defined a genomic distance based on SNVs. Firstly, we extracted the SNV sites (con_base) from each sample result file to obtain information about the reference base and its corresponding mutation type. We only considered the base that was supported by the most reads (con_base). Then, we merged all samples into a matrix where each row represented a SNV and each column represented the mutation type for each sample. We treated each column as a single sequence and aligned them to generate a phylogenetic distance matrix that contained the pairwise nucleotide substitution rate between samples using the Kimura 2-parameter method from the EMBOSS package [54]. To identify distinct strain clusters within species, the SNP haplotype distance matrix was normalized by dividing the maximal distance and hierarchical clustering was performed using the complete method from the R basic function hcluster. Kruskal test was used to access the genomic dissimilates.

### Protein 3D structure and functional prediction

The functions of some genes affected by SNVs were unknown. To address this, we used the AlphaFold2 artificial intelligence algorithm through ColabFold [55] and MMseqs2 [56] to model the protein structures based on multiple sequence alignments. Next, we predicted the functions of these proteins using DeepFRI [32], which is a Graph Convolutional Network that leverages sequence features from a protein language model and protein structures to predict protein functions.

### Association analysis

For each base site, there are four possible scenarios (i.e. A, T, C, G) for all samples regardless of the reference base. Initially, we excluded sites where a single base represented more than 90% of the population, and then we filtered out SNV sites that were present in less than 20% of the samples. Next, we only considered cases where two bases existed, which accounted for 97.36% of the data, and treated it as a binary variable. We then used linear and logistic regression models for continuous and binary traits, respectively, to establish microbial SNV associations with host phenotypes and metabolites. The formula used was: Metabolite/Phenotype ∼ SNV + day. To identify the effect of RMT on SNV, we used a logistic model with the following formula: SNV ∼ group + day.

### Ethical Approval

The study was approved by the institutional ethics review board of Hebei Agricultural University (ref. YS19003).

### Availability of data

All relevant data supporting the key findings of this study are available within the article and its Supplementary Information files. The raw metagenomic sequencing data used for the analysis presented in this study are available from the European Nucleotide Archive (ENA) under accession id PRJEB42631.

## Funding

This project was funded by the National Natural Science Foundation of China (NSFC) excellent young scientists fund program (overseas); the NSFC surface grant (32270077); the medical expert grant of Jiangsu; the Natural Science Foundation of Jiangsu grant (BK20220709); the Shuang Chuang project of Jiangsu (JSSCBS20221815), the NJMU starting grant (303073572NC21 & YNRCZN0301), the China agriculture research system of MOF and MARA project, and the Hebei dairy cattle innovation team of modern agro-industry technology research system (HBCT2018120203). The funders had no role in the study design, data collection and analysis, decision to publish, or preparation of the manuscript.

## Acknowledgements

We thank the management staffs of the study for their supports.

## Authors’ contributions

Y.S., L.T., X.K. and L.C. conceptualized and managed the study. Y.S., Y.L., T.W., Q.D., Y.G., Y.C., Q.L., J.S., H.Z., L.D., S.H., J.L., Y.G., Y.W., and L.C. collected the samples and generated the data. Q.D. and L.C. analysed the data. Q.D., D.H., X.W., Y.J. and L.C. drafted the manuscript. Q.D., D.H., X.W., Y.J., L.L., Q.D., T.W., H.Z., L.D., S.H., J.S., Y.W., H.Z., Y.S., W.S., Y.S., L.T., X.K. and L.C. reviewed and edited the manuscript.

## Consent for publication

All authors read and approved the final manuscript.

## Competing interests

The authors declare no competing interests.

## Abbreviations

SNVs: single nucleotide variations
MAGs: metagenomic assembled genomes
MT: microbial transplantation
SCFA: short-chain fatty acids
COG: clusters of orthologous genes
BCAA: branched-chain amino acid

## Notes

### Competing Interest Statement

The authors have declared no competing interest.

## References

1. Schwarzer M, Makki K, Storelli G, Machuca-Gayet I, Srutkova D, Hermanova P, Martino ME, Balmand S, Hudcovic T, Heddi A, et al: Lactobacillus plantarum strain maintains growth of infant mice during chronic undernutrition. Science 2016, 351:854-857.

2. Clavel T, Gomes-Neto JC, Lagkouvardos I, Ramer-Tait AE: Deciphering interactions between the gut microbiota and the immune system via microbial cultivation and minimal microbiomes. Immunol Rev 2017, 279:8-22.

3. Malmuthuge N, Guan LL: Understanding the gut microbiome of dairy calves: Opportunities to improve early-life gut health. J Dairy Sci 2017, 100:5996-6005.

4. Schwaiger K, Storch J, Bauer C, Bauer J: Development of selected bacterial groups of the rectal microbiota of healthy calves during the first week postpartum. J Appl Microbiol 2020, 128:366-375.

5. Takino T, Kato-Mori Y, Motooka D, Nakamura S, Iida T, Hagiwara K: Postnatal changes in the relative abundance of intestinal Lactobacillus spp. in newborn calves. J Vet Med Sci 2017, 79:452-455.

6. Song Y, Malmuthuge N, Steele MA, Guan LL: Shift of hindgut microbiota and microbial short chain fatty acids profiles in dairy calves from birth to pre-weaning. FEMS Microbiol Ecol 2018, 94.

7. Haley BJ, Kim SW, Salaheen S, Hovingh E, Van Kessel JAS: Differences in the Microbial Community and Resistome Structures of Feces from Preweaned Calves and Lactating Dairy Cows in Commercial Dairy Herds. Foodborne Pathog Dis 2020, 17:494-503.

8. Greenblum S, Carr R, Borenstein E: Extensive strain-level copy-number variation across human gut microbiome species. Cell 2015, 160:583-594.

9. Schloissnig S, Arumugam M, Sunagawa S, Mitreva M, Tap J, Zhu A, Waller A, Mende DR, Kultima JR, Martin J, et al: Genomic variation landscape of the human gut microbiome. Nature 2013, 493:45-50.

10. Sokurenko EV, Chesnokova V, Dykhuizen DE, Ofek I, Wu XR, Krogfelt KA, Struve C, Schembri MA, Hasty DL: Pathogenic adaptation of Escherichia coli by natural variation of the FimH adhesin. Proc Natl Acad Sci U S A 1998, 95:8922-8926.

11. Han B, Sivaramakrishnan P, Lin CJ, Neve IAA, He J, Tay LWR, Sowa JN, Sizovs A, Du G, Wang J, et al: Microbial Genetic Composition Tunes Host Longevity. Cell 2018, 173:1058.

12. Zeevi D, Korem T, Godneva A, Bar N, Kurilshikov A, Lotan-Pompan M, Weinberger A, Fu J, Wijmenga C, Zhernakova A, Segal E: Structural variation in the gut microbiome associates with host health. Nature 2019, 568:43-48.

13. Pasolli E, Asnicar F, Manara S, Zolfo M, Karcher N, Armanini F, Beghini F, Manghi P, Tett A, Ghensi P, et al: Extensive Unexplored Human Microbiome Diversity Revealed by Over 150,000 Genomes from Metagenomes Spanning Age, Geography, and Lifestyle. Cell 2019, 176:649-662 e620.

14. Bowers RM, Kyrpides NC, Stepanauskas R, Harmon-Smith M, Doud D, Reddy TBK, Schulz F, Jarett J, Rivers AR, Eloe-Fadrosh EA, et al: Minimum information about a single amplified genome (MISAG) and a metagenome-assembled genome (MIMAG) of bacteria and archaea. Nat Biotechnol 2017, 35:725-731.

15. Chaumeil PA, Mussig AJ, Hugenholtz P, Parks DH: GTDB-Tk v2: memory friendly classification with the Genome Taxonomy Database. Bioinformatics 2022.

16. Stewart RD, Auffret MD, Warr A, Wiser AH, Press MO, Langford KW, Liachko I, Snelling TJ, Dewhurst RJ, Walker AW, et al: Assembly of 913 microbial genomes from metagenomic sequencing of the cow rumen. Nat Commun 2018, 9:870.

17. Stewart RD, Auffret MD, Warr A, Walker AW, Roehe R, Watson M: Compendium of 4,941 rumen metagenome-assembled genomes for rumen microbiome biology and enzyme discovery. Nat Biotechnol 2019, 37:953-961.

18. Wilkinson T, Korir D, Ogugo M, Stewart RD, Watson M, Paxton E, Goopy J, Robert C: 1200 high-quality metagenome-assembled genomes from the rumen of African cattle and their relevance in the context of sub-optimal feeding. Genome Biol 2020, 21:229.

19. Seemann T: Prokka: rapid prokaryotic genome annotation. Bioinformatics 2014, 30:2068-2069.

20. Almeida A, Nayfach S, Boland M, Strozzi F, Beracochea M, Shi ZJ, Pollard KS, Sakharova E, Parks DH, Hugenholtz P, et al: A unified catalog of 204,938 reference genomes from the human gut microbiome. Nat Biotechnol 2021, 39:105-114.

21. Olm MR, Crits-Christoph A, Bouma-Gregson K, Firek BA, Morowitz MJ, Banfield JF: inStrain profiles population microdiversity from metagenomic data and sensitively detects shared microbial strains. Nat Biotechnol 2021, 39:727-736.

22. Wu S, Tian P, Tan T: Genomic landscapes of bacterial transposons and their applications in strain improvement. Appl Microbiol Biotechnol 2022, 106:6383-6396.

23. Müller N, Worm P, Schink B, Stams AJ, Plugge CM: Syntrophic butyrate and propionate oxidation processes: from genomes to reaction mechanisms. Environmental microbiology reports 2010, 2:489-499.

24. Galperin MY, Wolf YI, Makarova KS, Vera Alvarez R, Landsman D, Koonin EV: COG database update: focus on microbial diversity, model organisms, and widespread pathogens. Nucleic Acids Res 2021, 49:D274-D281.

25. Ku JWK, Gan YH: New roles for glutathione: Modulators of bacterial virulence and pathogenesis. Redox Biol 2021, 44:102012.

26. Wu H, Guang C, Zhang W, Mu W: Recent development of phenyllactic acid: physicochemical properties, biotechnological production strategies and applications. Crit Rev Biotechnol 2021:1-16.

27. Malik N, Kriplani A, Agarwal N, Bhatla N, Kachhawa G, Yadav RK: Dehydroepiandrosterone as an adjunct to gonadotropins in infertile Indian women with premature ovarian aging: A pilot study. J Hum Reprod Sci 2015, 8:135-141.

28. Landete JM, Arena ME, Pardo I, Manca de Nadra MC, Ferrer S: The role of two families of bacterial enzymes in putrescine synthesis from agmatine via agmatine deiminase. Int Microbiol 2010, 13:169-177.

29. Lefevre PL, Palin MF, Murphy BD: Polyamines on the reproductive landscape. Endocr Rev 2011, 32:694-712.

30. Labarrere CA, Kassab GS: Glutathione: A Samsonian life-sustaining small molecule that protects against oxidative stress, ageing and damaging inflammation. Front Nutr 2022, 9:1007816.

31. Jumper J, Evans R, Pritzel A, Green T, Figurnov M, Ronneberger O, Tunyasuvunakool K, Bates R, Zidek A, Potapenko A, et al: Highly accurate protein structure prediction with AlphaFold. Nature 2021, 596:583-589.

32. Gligorijevic V, Renfrew PD, Kosciolek T, Leman JK, Berenberg D, Vatanen T, Chandler C, Taylor BC, Fisk IM, Vlamakis H, et al: Structure-based protein function prediction using graph convolutional networks. Nat Commun 2021, 12:3168.

33. Liao X, Deng T, Zhu Y, Du G, Chen J: Enhancement of glutathione production by altering adenosine metabolism of Escherichia coli in a coupled ATP regeneration system with Saccharomyces cerevisiae. J Appl Microbiol 2008, 104:345-352.

34. Ballatori N, Hammond CL, Cunningham JB, Krance SM, Marchan R: Molecular mechanisms of reduced glutathione transport: role of the MRP/CFTR/ABCC and OATP/SLC21A families of membrane proteins. Toxicol Appl Pharmacol 2005, 204:238-255.

35. Raper HS: The Tyrosinase-tyrosine Reaction: Production from Tyrosine of 5: 6-Dihydroxyindole and 5: 6-Dihydroxyindole-2-carboxylic Acid-the Precursors of Melanin. Biochem J 1927, 21:89-96.

36. Shuster V, Fishman A: Isolation, cloning and characterization of a tyrosinase with improved activity in organic solvents from Bacillus megaterium. J Mol Microbiol Biotechnol 2009, 17:188-200.

37. Yassour M, Jason E, Hogstrom LJ, Arthur TD, Tripathi S, Siljander H, Selvenius J, Oikarinen S, Hyoty H, Virtanen SM, et al: Strain-Level Analysis of Mother-to-Child Bacterial Transmission during the First Few Months of Life. Cell Host Microbe 2018, 24:146-154 e144.

38. Vatanen T, Plichta DR, Somani J, Munch PC, Arthur TD, Hall AB, Rudolf S, Oakeley EJ, Ke X, Young RA, et al: Genomic variation and strain-specific functional adaptation in the human gut microbiome during early life. Nat Microbiol 2019, 4:470-479.

39. Li Y, Shen Y, Niu J, Guo Y, Pauline M, Zhao X, Li Q, Cao Y, Bi C, Zhang X, et al: Effect of active dry yeast on lactation performance, methane production, and ruminal fermentation patterns in early-lactating Holstein cows. J Dairy Sci 2021, 104:381-390.

40. Van Soest PJ, Robertson JB, Lewis BA: Methods for dietary fiber, neutral detergent fiber, and nonstarch polysaccharides in relation to animal nutrition. J Dairy Sci 1991, 74:3583-3597.

41. Rice EM, Aragona KM, Moreland SC, Erickson PS: Supplementation of sodium butyrate to postweaned heifer diets: Effects on growth performance, nutrient digestibility, and health. J Dairy Sci 2019, 102:3121-3130.

42. Langmead B, Trapnell C, Pop M, Salzberg SL: Ultrafast and memory-efficient alignment of short DNA sequences to the human genome. Genome Biol 2009, 10:R25.

43. Langmead B, Wilks C, Antonescu V, Charles R: Scaling read aligners to hundreds of threads on general-purpose processors. Bioinformatics 2019, 35:421-432.

44. Bolger AM, Lohse M, Usadel B: Trimmomatic: a flexible trimmer for Illumina sequence data. Bioinformatics 2014, 30:2114-2120.

45. Li D, Liu CM, Luo R, Sadakane K, Lam TW: MEGAHIT: an ultra-fast single-node solution for large and complex metagenomics assembly via succinct de Bruijn graph. Bioinformatics 2015, 31:1674-1676.

46. Li H, Handsaker B, Wysoker A, Fennell T, Ruan J, Homer N, Marth G, Abecasis G, Durbin R, Genome Project Data Processing S: The Sequence Alignment/Map format and SAMtools. Bioinformatics 2009, 25:2078-2079.

47. Kang DD, Li F, Kirton E, Thomas A, Egan R, An H, Wang Z: MetaBAT 2: an adaptive binning algorithm for robust and efficient genome reconstruction from metagenome assemblies. PeerJ 2019,7:e7359.

48. Parks DH, Imelfort M, Skennerton CT, Hugenholtz P, Tyson GW: CheckM: assessing the quality of microbial genomes recovered from isolates, single cells, and metagenomes. Genome Res 2015, 25:1043-1055.

49. Olm MR, Brown CT, Brooks B, Banfield JF: dRep: a tool for fast and accurate genomic comparisons that enables improved genome recovery from metagenomes through de-replication. ISME J 2017, 11:2864-2868.

50. Olm MR, Crits-Christoph A, Diamond S, Lavy A, Matheus Carnevali PB, Banfield JF: Consistent Metagenome-Derived Metrics Verify and Delineate Bacterial Species Boundaries. mSystems 2020, 5.

51. Parks DH, Chuvochina M, Waite DW, Rinke C, Skarshewski A, Chaumeil PA, Hugenholtz P: A standardized bacterial taxonomy based on genome phylogeny substantially revises the tree of life. Nat Biotechnol 2018, 36:996-1004.

52. Letunic I, Bork P: Interactive Tree Of Life (iTOL) v4: recent updates and new developments. Nucleic Acids Res 2019, 47:W256-W259.

53. Hyatt D, Chen GL, Locascio PF, Land ML, Larimer FW, Hauser LJ: Prodigal: prokaryotic gene recognition and translation initiation site identification. BMC Bioinformatics 2010, 11:119.

54. Rice P, Longden I, Bleasby A: EMBOSS: the European Molecular Biology Open Software Suite. Trends Genet 2000, 16:276-277.

55. Mirdita M, Schutze K, Moriwaki Y, Heo L, Ovchinnikov S, Steinegger M: ColabFold: making protein folding accessible to all. Nat Methods 2022, 19:679-682.

56. Steinegger M, Soding J: MMseqs2 enables sensitive protein sequence searching for the analysis of massive data sets. Nat Biotechnol 2017, 35:1026-1028.

